# Recombination rate variation shapes genomic variability of phylogeographic structure in a widespread North American songbird (Aves: *Certhia americana*)

**DOI:** 10.1101/2022.09.25.509431

**Authors:** Joseph D. Manthey, Garth M. Spellman

## Abstract

The nonrandom distribution of chromosomal characteristics and functional elements—genomic architecture—impacts the relative strengths and impacts of population genetic processes across the genome. Due to this relationship, genomic architecture has the potential to shape variation in population genetic structure across the genome. Population genetic structure has been shown to vary across the genome in a variety of taxa, but this body of work has largely focused on pairwise population genomic comparisons between closely related taxa. Here, we used whole genome sequencing of seven phylogeographically structured populations of a North American songbird, the Brown Creeper (*Certhia americana*), to determine the impacts of genomic architecture on phylogeographic structure variation across the genome. Using multiple methods to infer phylogeographic structure—ordination, clustering, and phylogenetic methods— we found that recombination rate variation explained a large proportion of phylogeographic structure variation. Genomic regions with low recombination showed phylogeographic structure consistent with the genome-wide pattern. In regions with high recombination, we found strong phylogeographic structure, but with discordant patterns relative to the genome-wide pattern. In regions with high recombination rate, we found that populations with small effective population sizes evolve relatively more rapidly than larger populations, leading to discordant signatures of phylogeographic structure. These results suggest that the interplay between recombination rate variation and effective population sizes shape the relative impacts of linked selection and genetic drift in different parts of the genome. Overall, the combined interactions of population genetic processes, genomic architecture, and effective population sizes shape patterns of variability in phylogeographic structure across the genome of the Brown Creeper.

## INTRODUCTION

The formation of population structure is implicit in the speciation process (Allmon, 1992; Mayr et al., 1963). Speciation is a continuum; early differentiating populations may have few genomic regions with evidence for genetic differentiation or barriers to gene flow (Ellegren et al., 2012; Nosil and Feder, 2012; Toews et al., 2016; Wu, 2001) while later in the speciation process two diverging populations may exhibit strong genetic differentiation across their entire genomes (Nosil and Feder, 2012; Ravinet et al., 2018; Wu, 2001). Across the genome, variation in local evolutionary history is shaped by the relative strengths of the population genomic processes of natural selection, gene flow, and genetic drift. For example, population genetic structure may differ in genomic regions with strong natural selection or differential patterns of gene flow relative to the rest of the genome (Louis et al., 2020; Whiting et al., 2021).

Variable impacts of selection, drift, and gene flow across the genome may be partially attributed to genomic architecture—the nonrandom distribution of chromosomal characteristics and functional elements (Koonin, 2009)—whereby variation in gene content, recombination rates, or other genomic characteristics impact the relative effects of population genomic processes. For example, we expect that genomic regions with lower recombination rates will be relatively depleted of genetic diversity and genetic differentiation will accumulate faster, due to relatively increased effects of linked selection (Cruickshank and Hahn, 2014; Haenel et al., 2018; Hey and Kliman, 2002; Nachman and Payseur, 2012). Indeed, speciation and population genomic studies have often found a negative relationship between recombination rates and genetic differentiation across the genome (Kulathinal et al., 2008; Manthey et al., 2021; Roesti et al., 2013; Stankowski et al., 2019; Vijay et al., 2016).

If genomic architecture impacts population genomic patterns within populations and between pairs of taxa, we may naturally extend this idea to phylogeographic patterns in taxa with multiple lineages at different stages of evolutionary distinctiveness. As such, we may predict that estimates of phylogeographic structure will vary across the genome and be shaped by several factors and their interactions: (1) population size variation of differentiating populations shaping relative strengths of selection and drift, (2) time since divergence began, (3) genomic architecture, and (4) quantity of interpopulation gene flow. Exemplar taxa to better understand the impacts of genomic architecture on population genetic structure are species (or species complexes) with multiple lineages on distinct evolutionary trajectories, populations with different effective sizes, and genomes exhibiting heterogeneous genomic architecture.

The Brown Creeper (*Certhia americana*) is an excellent study organism to understand the impacts of genomic architecture on phylogeographic structure; its genome is highly heterogeneous in both structure and content, and it has seven distinct phylogeographic lineages (Manthey et al., 2011a; Manthey et al., 2015). The genome of the Brown Creeper, as like other birds, has chromosomes that (1) span two orders of magnitude in size, (2) vary in their gene and repetitive element content, and (3) exhibit local effective recombination rates that can vary by an order of magnitude or more (Dutoit et al., 2017; Ellegren, 2010; Kapusta and Suh, 2017; Kawakami et al., 2014; Manthey et al., 2021). Additionally, the Brown Creeper has strong phylogeographic structure, with two main lineages that have been diverging around one million years (Fig. 1A), multiple phylogeographically structured clades within each of these two main lineages, and populations with different effective sizes (Manthey et al., 2011a, b; Manthey et al., 2015, 2021).

**Figure 1.**
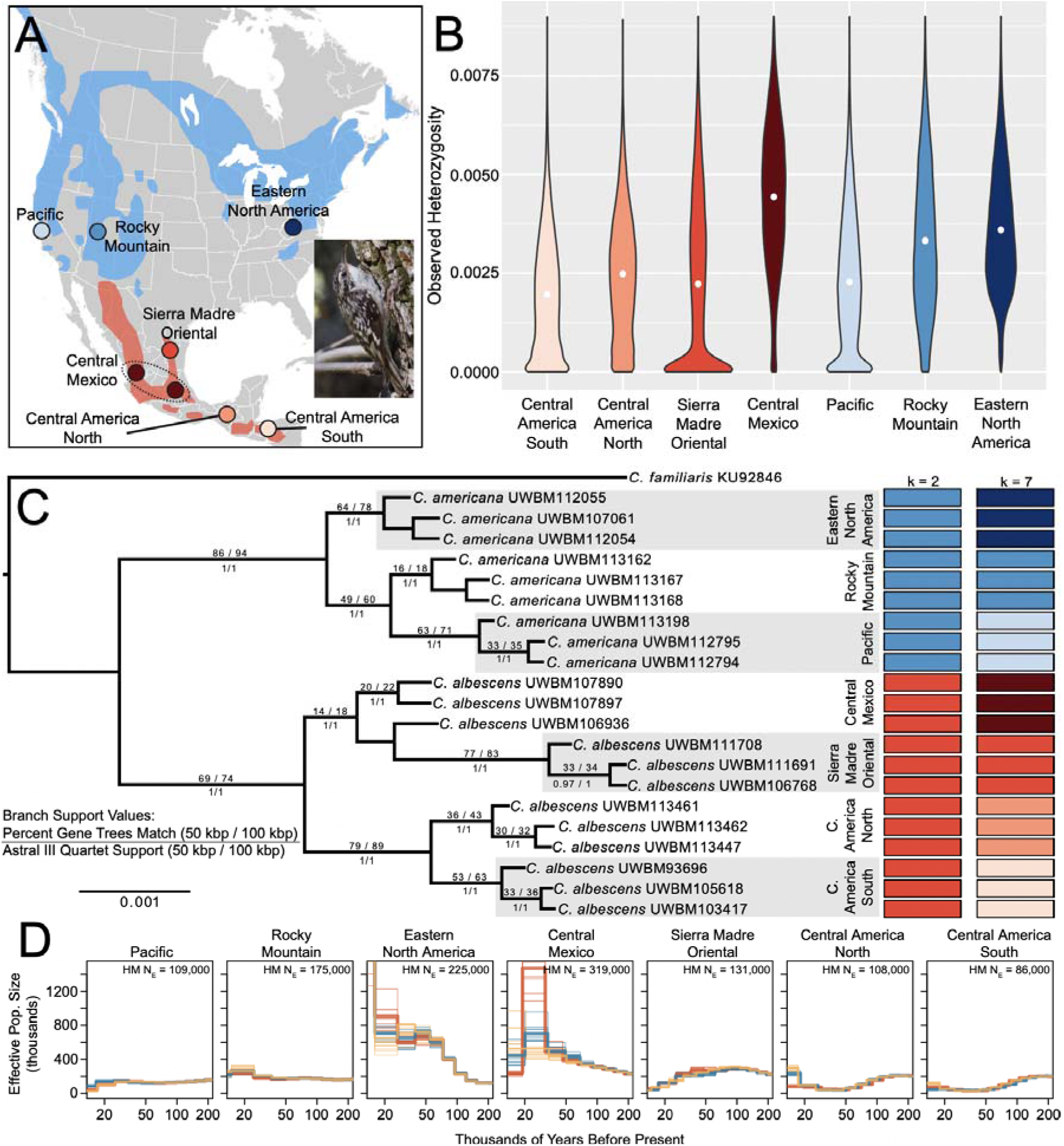
Sampling map, genetic diversity, phylogeographic structure, and demography. (A) Sampling localities for this study. Colors for localities are consistent across plots. (B) Violin plots of observed heterozygosity across all 50 kbp sliding windows. White circles indicate means. (C) Consensus phylogeny using 19,639 gene trees from the 50 kbp windows. Support values are labelled for all branches showing the highest support from all estimation methods (N = 4), including consensus and ASTRAL phylogenies for both the 50 kbp and 100 kbp window datasets. At tips of phylogeny are ADMIXTURE results from 19,602 genome-wide SNPs (thinned every 50 kbp) for either two or seven assumed genetic clusters (k = 2 or k = 7, respectively). (D) Demographic history estimated with MSMC2. Different individuals are represented with differently colored lines and bootstraps are shown with thin lines. In (D), HM = harmonic mean. Brown Creeper photo in (A) by JDM from the Chiricahua Mountains in southern Arizona.

Using the Brown Creeper as our study taxon, we used whole-genome sequencing data to assess how genome architecture influences phylogeographic structure; we used phylogenetic, ordination, and population genetic clustering methods to characterize phylogeographic structure across the genome. Our aims here were twofold. First, we looked to decipher how genomic architecture and effective population sizes shape phylogeographic structure variation across the genome. Second, because phylogeographic structure may be estimated in many ways, we wanted to identify whether phylogeographic signal variation was consistent using different methodologies.

Based on previous population genomic research, we hypothesized that genomic regions with low recombination rates will have faster lineage sorting. Based on this hypothesis, we would predict that phylogeographic estimates consistent with the species or population level evolutionary history of the group will be concentrated in genomic regions with low recombination. Additionally, we may predict a couple alternative patterns in high recombination regions. We may identify mixed or weak phylogeographic structure in high recombination regions. Alternatively, we may identify strong, but divergent patterns of phylogeographic structure in high recombination regions linked with varied molecular evolutionary rates among sampled populations. We may also predict an interaction between recombination rate and population sizes; whereby, relatively smaller populations will undergo faster molecular evolution in genomic regions characterized by stabilizing selection, such as in gene-dense, high recombination genomic regions.

## MATERIALS AND METHODS

### Study organism, sampling, and sequencing

The Brown Creeper is a songbird found in forested regions of the Americas; it has a widespread distribution ranging from Honduras in the south to Alaska in the north (Fig. 1A) (Poulin et al., 2020). The Brown Creeper is aptly named; it creeps up trees in search of its largely invertebrate diet and has dark coloration on its back and sides (Fig. 1A inset) that makes for excellent camouflage on trees with dark bark. In previous work using tens to thousands of genetic markers, we have identified phylogeographic structure in the Brown Creeper, with up to seven distinct lineages (Manthey et al., 2011a, b; Manthey et al., 2015). Here, we aimed to obtain population genomic sequencing data for seven populations with three individuals sampled per population (Fig 1, Table S1). We downloaded previously published sequencing data from two of these populations and one outgroup sample of *C. familiaris* (Table S1) (Manthey et al., 2021). For the other five populations, we obtained representative tissue samples from natural history museums and extracted genomic DNA using QIAGEN DNeasy blood and tissue kits following manufacturer guidelines. DNA from extractions was used to create standard Illumina sequencing libraries, followed by sequencing on an Illumina NovaSeq6000 at the Texas Tech University Center for Biotechnology and Genomics with the goal of obtaining ∼15-30× coverage per individual.

### Reference genome and genomic architecture

For the reference genome, we used the chromosome-scale *Certhia americana* genome we assembled for a prior study (NCBI assembly ASM1869719v1; Manthey et al. 2021). From data reported in this previous study (Manthey et al. 2021), we summarized mean effective recombination rates for each of the two main Brown Creeper lineages, transposable element (TE) and gene density, and GC content across the genome in 50kbp and 100 kbp non-overlapping sliding windows (Fig. 2). We used the mean effective recombination rates of the two lineages because they were highly correlated (r = 0.829-0.861 across different correlation metrics; all p << 0.001), and we can assume relatively conserved recombination rates over short evolutionary time frames (Singhal et al., 2015).

**Figure 2.**
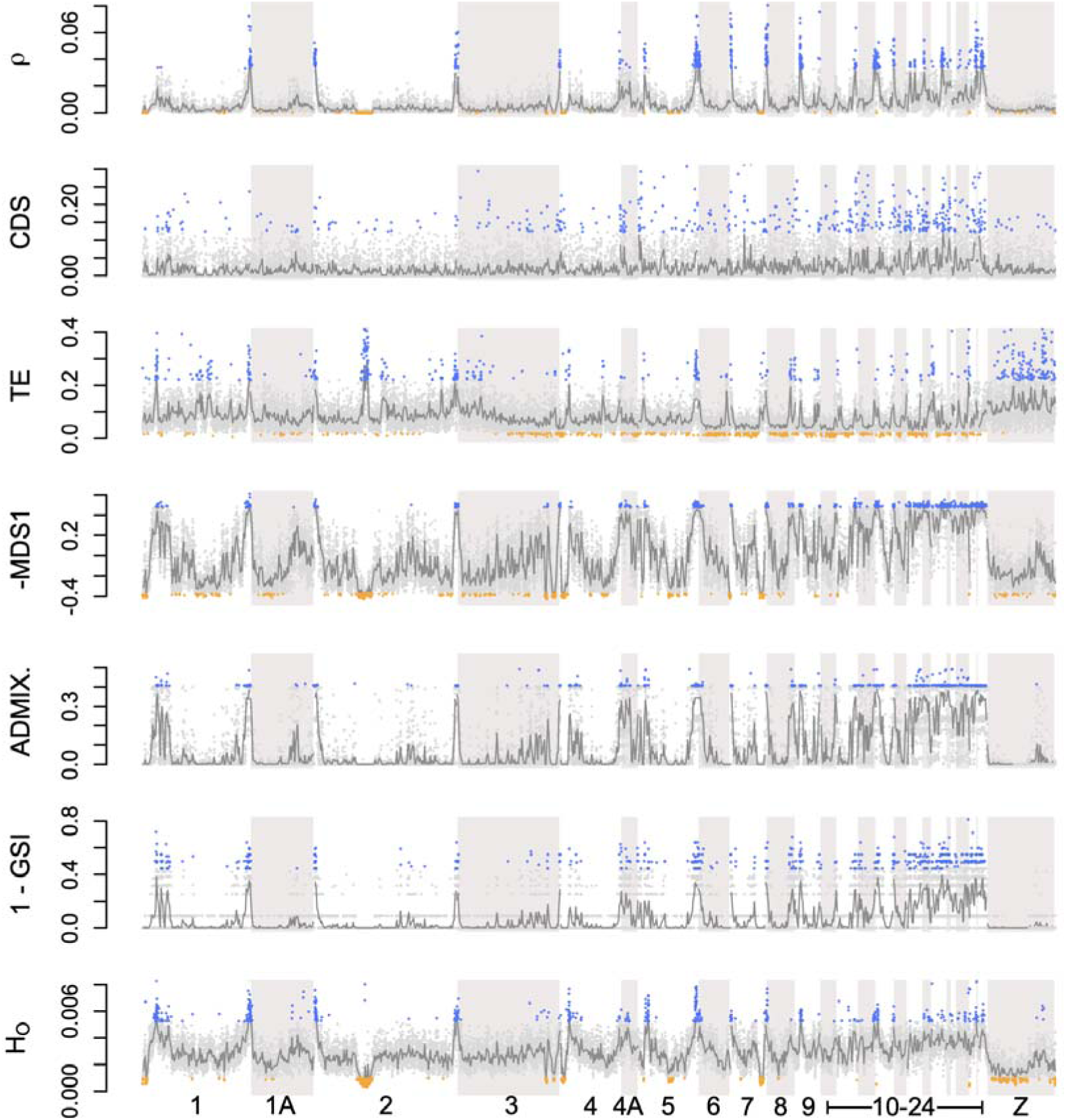
Genomic architecture and phylogeographic structure variation across the genome in 50kbp sliding windows. Gray points indicate window values and dark gray lines indicate means across 20 windows (i.e., 1 Mbp windows). Blue indicates top 2.5% of outliers and orange indicates bottom 2.5% of outliers for each statistic. Bottom outliers not shown for CDS, ADMIX., and GSI as these each have large proportions of values equal to zero. Abbreviations: mean effective recombination rate of the two main lineages (ρ), gene content (CDS), transposable element and repetitive DNA content (TE), multidimensional scaling axis one from principal components analyses (-MDS1), mean genealogical sorting index for the two main *Certhia* lineages (1 – GSI), ADMIXTURE deviation relative to genome-wide analysis (ADMIX.), and mean genetic diversity across individuals as measured using observed heterozygosity (H_O_). MDS1 is plotted as negative and GSI is plotted as (1 – GSI) so that deviations from expected phylogeographic structure for MDS1, GSI, and ADMIXTURE are all represented as higher values.

Briefly, these estimates were initially obtained with the following methods: effective recombination rate was estimated using LDhat (McVean and Auton, 2007), TEs were annotated with RepeatMasker v4.08 (Smit et al., 2015), and gene content was annotated with MAKER v2.31.10 (Cantarel et al., 2008).

### Sequencing data filtering and genotyping

We used the program bbduk (Bushnell, 2014) to quality filter raw sequencing data. We then use BWA v0.7.17 (Li and Durbin, 2009) to align filtered reads to the *Certhia americana* reference genome. Next, we used samtools v1.4.1 (Li et al., 2009) to convert the BWA output SAM file to BAM format, followed by using the Genome Analysis Toolkit (GATK) v4.1.0.0 (McKenna et al., 2010) to clean, sort, add read groups to, and remove duplicates from BAM files. We used the samtools ‘depth’ command to measure sequencing coverage across the reference genome for each individual (Fig. S1, Table S1).

We then used GATK in three steps to genotype all individuals: we used the ‘HaplotypeCaller’ function to call preliminary genotypes for each individual, built a database for all samples using the ‘GenomicsDBImport’ function, and lastly used the ‘GenotypeGVCFs’ function to group genotype all individuals together for both variant and invariant sites. We filtered the genotype data using VCFtools v0.1.14 (Danecek et al., 2011) with the following restrictions: (1) minimum site quality of 20, (2) minimum genotype quality of 20, (3) minimum depth of coverage of 6, (4) maximum mean depth of coverage of 50, and (5) removal of indels. For downstream analyses, we used various settings for (1) amount of missing data allowed, (2) whether the outgroup was included, (3) inclusion or exclusion of invariant sites, and (4) thinning between sites (see Table 1 for exact dataset characteristics used for all seven datasets used for analyses). Although it has been suggested that variation in minor allele frequency (MAF) filtering may impact estimates of genetic structure (Linck and Battey, 2019), we did not filter for a minimum MAF because we previously showed that changing MAF did not significantly impact pairwise F_ST_ calculations in *C. americana* (Manthey et al., 2021).

**Table 1.** Different datasets used in this study and their characteristics. For inclusion, all datasets required sites to have a minimum quality score of 20, minimum of 20 genotype quality, a minimum depth per individual for a site to be called of 6, and a max mean depth across individuals of 50.

| Set | Missing Allowed | OG<sup>1</sup> | Window size | # Windows | Min. Sep. per SNP<sup>2</sup> | # SNPs | # Sites | Analyses |
| --- | --- | --- | --- | --- | --- | --- | --- | --- |
| A | 0% |  | - | - | 50 kbp | 19,602 | 19,602 | Genome-wide ADMIXTURE |
| B | 0% |  | - | - | 100 kbp | 9,838 | 9,838 | Genome-wide ADMIXTURE |
| C | 20% | X | 50 kbp | 19,713 | - | 35,463,408 | 890,335,872 | RAxML, GSI, H<sub>o</sub> |
| D | 20% | X | 100 kbp | 9,862 | - | 35,463,408 | 890,335,872 | RaxML, GSI, H<sub>o</sub> |
| E | 0% |  | - | - | - | 21,657,220 | 21,657,220 | LOSTRUCT |
| F | 0% |  | 50 kbp | 19,697 | - | 21,657,220 | 21,657,220 | ADMIXTURE per window |
| G | 0% |  | 100 kbp | 9,860 | - | 21,657,220 | 21,657,220 | ADMIXTURE per window |
| H | 0% |  | - | - | - | * | * | MSMC2 |

<sup>1</sup>OG = outgroup included
<sup>2</sup>Min. Sep. per SNP = minimum separation between SNPs (i.e., thinning)
\*SNPs and sites not labeled for MSMC2 analyses because these values varied per individual. For each individual, no missing data was allowed for that particular individual's MSMC2 demographic analyses input files. The MSMC2 dataset did not include the Z chromosome.

### Phylogenomics

We estimated “gene trees” in non-overlapping windows (window size 50 kbp and 100 kbp; datasets C and D in Table 1) using RaxML v8.2.12 (Stamatakis, 2014) with the GTRCAT model of sequence evolution. For a window to be included, we required a minimum of 10 kbp in the alignment following filtering. This resulted in 19,639 phylogenies for the 50 kbp sliding windows and 9846 phylogenies for the 100 kbp sliding windows. We summarized the best-supported trees using the sumtrees.py script, part of the DendroPy Python package (Sukumaran and Holder, 2010). Lastly, we used ASTRAL III v 5.6.3 (Zhang et al., 2018) to calculate a species tree from all gene trees, using quartet frequencies as a measure of local support (Sayyari and Mirarab, 2016).

For each gene tree, we estimated the genealogical sorting index (GSI) (Cummings et al., 2008) for (1) each of the seven populations, and (2) each of the two main lineages (i.e., north and south shaded blue and red, respectively, in Fig. 1). The GSI statistic uses rooted gene trees to quantify lineage sorting; the GSI quantifies exclusive ancestry of individuals in labelled groups by dividing the minimum number of nodes to produce monophyly for a defined group divided by the observed number of nodes necessary to connect all individuals of a group in a phylogeny. The maximum GSI value is one and occurs when a defined group is monophyletic. We calculated the GSI statistic for each sampled population and for each of the two main lineages (Fig. 1). We expect that each sampled population will show evidence of phylogenetic clustering, regardless of position in the genome. As such, GSI at the population level may have little variation tied to genomic architecture. In contrast, which populations group with which other populations are likely to be influenced by rates of lineage sorting and are therefore likely influenced by genomic architecture. As such, we may expect GSI measures at the lineage level (i.e., two main clades) to be impacted by genomic architecture.

To obtain a relative rate of molecular evolution for each population per genomic window, we calculated root-to-tip distances (RTD) in our phylogenies. For each individual across every gene tree, we calculated the RTD using the R package ape (Paradis et al., 2004). As a summary of these RTDs, we calculated the mean RTD per population relative to the maximum RTD value for each gene tree (i.e., this relative statistic’s range = 0 – 1). We then measured the relationships between harmonic mean effective population sizes (mean per population) and relative RTD in regions with low recombination and high recombination, as well as the shift in values between high and low recombination regions. We also estimated relationships between these characteristics while accounting for the evolutionary relationships of the samples using phylogenetic independent contrasts (PICs) calculated in the R package ape (Paradis et al., 2004).

### Genetic structure

We estimated genetic structure using the program ADMIXTURE v1.3.0 (Alexander et al., 2009). Here, we used genome-wide SNPs thinned to either a minimum of 50 kbp or 100 kbp distance between SNPs (datasets A and B in Table 1). We ran ADMIXTURE with an assumed number of genetic clusters equal to two or seven (i.e., k = 2 and k = 7) based on our knowledge of phylogeographic structure in this species from previous studies (Manthey et al., 2011a, b; Manthey et al., 2015). To assess how genetic structure estimation varied across the genome, we also ran ADMIXTURE for SNP datasets in non-overlapping sliding windows of 50 kbp and 100 kbp (datasets E and F in Table 1). We compared the genome-wide ADMIXTURE results to those in each window by summing the differences in group membership for each individual. In other words, this deviation value ranges from zero to nearly one, where a value of zero is identical group assignment for all individuals with the same admixture coefficients for each genetic cluster, and higher values indicate more dissimilar matrices.

We also estimated variation in genetic structure across the genome using LOSTRUCT (Li and Ralph, 2019). This method runs in three steps: (1) reducing the dimensionality of the data using principal component analysis (PCA) for each genomic window, (2) find distances between PCA maps, and (3) using multidimensional scaling (MDS) to display variation in PCA-based genetic structure across the genome. We used LOSTRUCT in sliding windows of 50 kbp and 100 kbp (datasets E and F in Table 1). We used variation in MDS axis one (hereafter MDS1) to describe variation in these PCA-based estimates of genetic structure across the genome. We expect that genomic regions with low recombination will exhibit strong phylogeographic structure similar to genome-wide patterns (e.g., Fig. 1). In contrast, we expect that genomic regions with high recombination will either exhibit little genetic structure or alternative patterns of genetic structure. As such, we expect that variation in ADMIXTURE and PCA results will covary with genomic architecture.

### Genetic diversity

Across 50 kbp and 100 kbp sliding windows (datasets C and D in Table 1), we measured observed heterozygosity per individual as a measure of genetic diversity.

### Population genomic correlations

We used the R package Hmisc (Harrell and Dupont, 2020) to estimate both Pearson and Spearman correlation coefficients between population genomic summary statistics and characteristics of genomic architecture in sliding windows. We used the R package corrplot (Wei et al., 2017) to visualize correlations between all summary statistics. Lastly, we used variance partitioning, implemented in the R package vegan (Oksanen et al., 2007), to assess the proportion of variance in phylogeographic structure estimates explained by genomic architecture.

### Population demographic history

We used the program MSMC2 v1.1.0 (Schiffels and Durbin, 2014) to estimate demographic history for each individual. For use in MSMC, we masked regions of the genome not genotyped due to low coverage or low genotype quality scores. Additionally, we did not include the sex chromosomes in demography calculations. We ran MSMC for each individual allowing up to 20 iterations (default setting) and used up to 23 inferred distinct time segments because this setting worked well to reduce spurious or inconsistent results in other songbirds that we have studied (Manthey et al., 2022). We performed 10 bootstraps for each demography estimate, using 1 Mbp bootstrapped segments of the genome, to assess how demographic signal varies when subsetting parts of the genome. Because the output of MSMC is interpreted relative to assumed mutation rates and generation times, we used the Brown Creeper genome mutation rate estimate of 2.506 × 10^-9^ substitutions per site per year (Manthey et al., 2021). Because there are no published estimates of Brown Creeper generation times, we used a proxy generation time of double the age of sexual maturity (Nadachowska-Brzyska et al., 2015) using maturity estimates from the Animal Aging and Longevity Database (age of maturity = one year) (Tacutu et al., 2017). With the MSMC demographic model output, we estimated the harmonic mean population sizes for each population by using the mean MSMC population size estimates for each individual in 1000-year discrete time intervals over the past 200 kya.

### Code availability

All computer code used for analyses and figure creation (applicable for some figures) for this project is available here: <u>github.com/jdmanthey/certhia_phylogeography</u>. We used the following R packages for figure creation and file manipulation that were not cited in other parts of the methods section: Biostrings (Pagès et al., 2017), ggplot2 (Wickham, 2011), palettetown (Lucas, 2016), phytools (Revell, 2012), and RcolorBrewer (Neuwirth, 2014).

## RESULTS

### Genome-wide phylogeographic structure and demography

We used genomic sequencing data from 21 ingroup individuals sequenced at ∼18-29× genomic coverage to estimate genome-wide patterns of phylogeographic structure. We corroborated previous genetic work (that used few to thousands of genetic markers) by identifying hierarchical and strong phylogeographic structure among sampled populations (Manthey et al., 2011a, b; Manthey et al., 2015). The deepest phylogenomic split separates northern from southern populations and there is additional support for distinctiveness of each of the regionally sampled populations (Fig. 1C).

Genome-wide ADMIXTURE results were consistent across different thinning strategies (i.e., 50 kbp and 100 kbp thinning; only 50 kbp results plotted) and showed patterns of genetic structure that aligned well with the species tree (Fig. 1C). Demographic history estimates for each individual were consistent within populations, but each population exhibited a distinct demographic history (Fig. 1D). In the Pacific and Central American localities, population sizes have fluctuated somewhat, but generally stay below an N_E_ of 200,000 over the past 100 ky (Fig. 1D). The Sierra Madre Oriental population showed evidence for a sharp decline in N_E_ over the past 50 ky, shifting from an N_E_ ∼ 200,000 at about 50 kya to an N_E_ ∼ 10,000 in the past ten kya (Fig. 1D). The Rocky Mountains population has remained relatively stable over the past 100 ky with an N_E_ ∼ 200,000 (Fig. 1D). Lastly, the Eastern North America and Central Mexico populations have exhibited fluctuations and much larger N_E_ than all other populations over the past 100 ky (Fig. 1D). Harmonic mean effective population sizes over the past 200 ky ranged from ∼85,000 (Central America South) to ∼320,000 (Central Mexico) (Fig. 1D). Harmonic mean effective population size estimates over the past 200 ky are highly correlated with observed heterozygosity estimates for each individual (r = 0.962).

### Genetic diversity

Genetic diversity in each individual varied widely across the genome, with mean observed heterozygosity values for each individual between ∼0.00165 and 0.00463 (Fig 1B). Observed heterozygosity was strongly correlated across the genome in all pairwise comparisons of individuals (mean r = 0.409, range = 0.189 – 0.853; all p << 0.001). Notably, the Sierra Madre Oriental, Pacific, and Central American localities had more windows with no heterozygosity than the other populations (Fig. 1B).

### Phylogeographic structure variation across the genome

We estimated phylogeographic structure across the genome using PCA, ADMIXTURE, and the GSI metric from phylogenies; all estimates of phylogeographic structure varied across the genome, with the outliers deviating the most from genome-wide patterns clustering mostly on small chromosomes and the ends of large chromosomes (Fig. 2; Fig. S2). The most extreme deviations from genome-wide phylogeographic structure appear to cluster the Central Mexico population with the northern lineage (Fig. S3), and in some cases also cluster the Sierra Madre Oriental population with the northern lineage (Fig. S3; Fig. 3). As an example, we plot the phylogeographic structure for one of these outlier windows in Fig. 3, demonstrating the clustering of the Sierra Madre Oriental and Central Mexico populations with the northern lineage.

**Figure 3.**
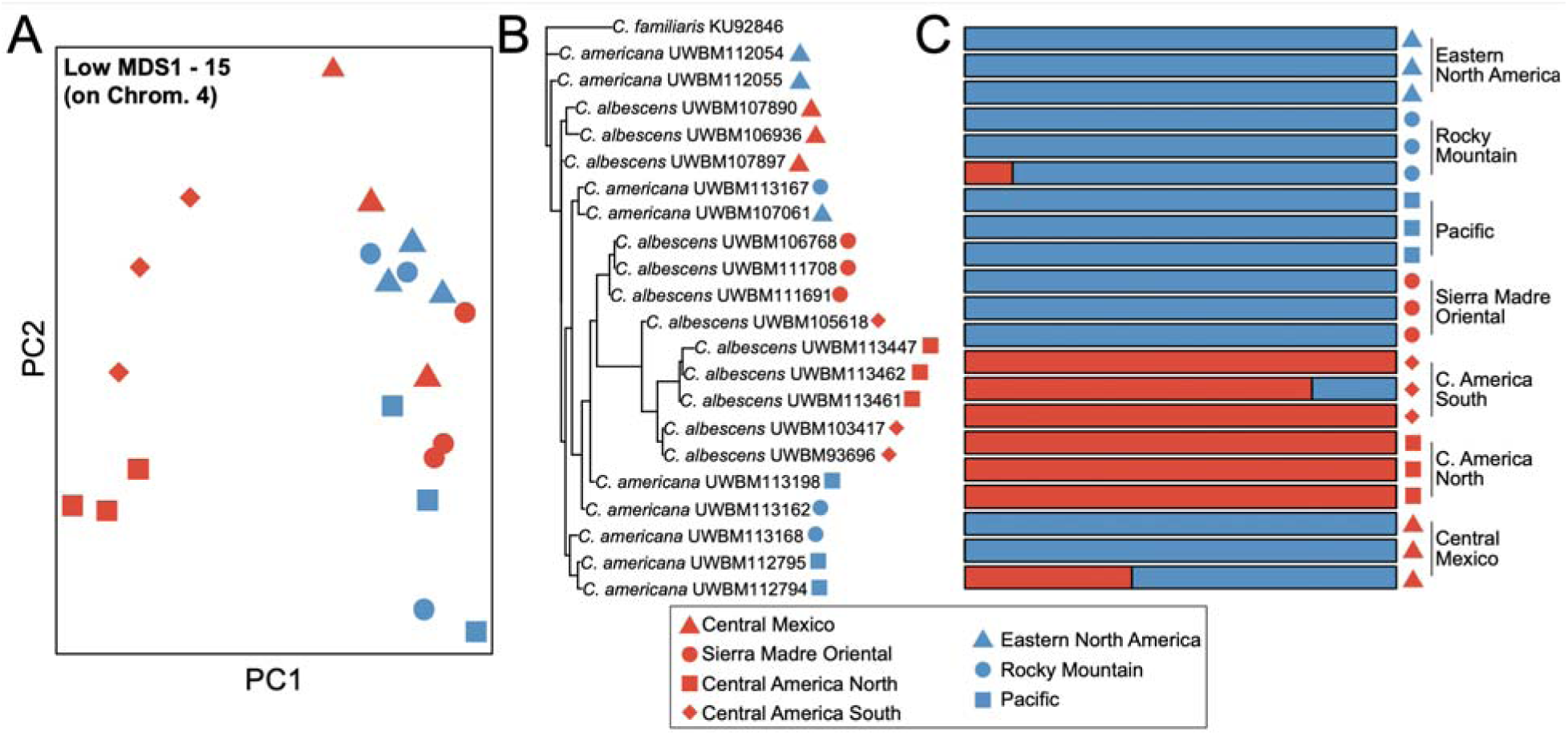
Example phylogeographic structure in a window that deviates from genome-wide patterns. This window had the 15^th^ lowest MDS1 value of all windows from the principal components analysis (PCA) across the genome. (A) PCA, (B) RAxML phylogeny, and (C) ADMIXTURE results for a window on chromosome 4.

### Correlations between phylogeographic structure estimates and genomic architecture

We found a strong correlation between different measures of phylogeographic structure, including GSI at the lineage level, MDS1 of the PCA, and ADMIXTURE deviations (all r ≥ 0.58; p <<< 0.001; Fig. 4; Fig. S4). In contrast, GSI at the population level did not vary consistently with the other metrics (all r between –0.21 and –0.03), likely because the population-level GSI usually showed similar patterns across the genome (i.e., individuals generally clustered with individuals sampled from the same population).

**Figure 4.**
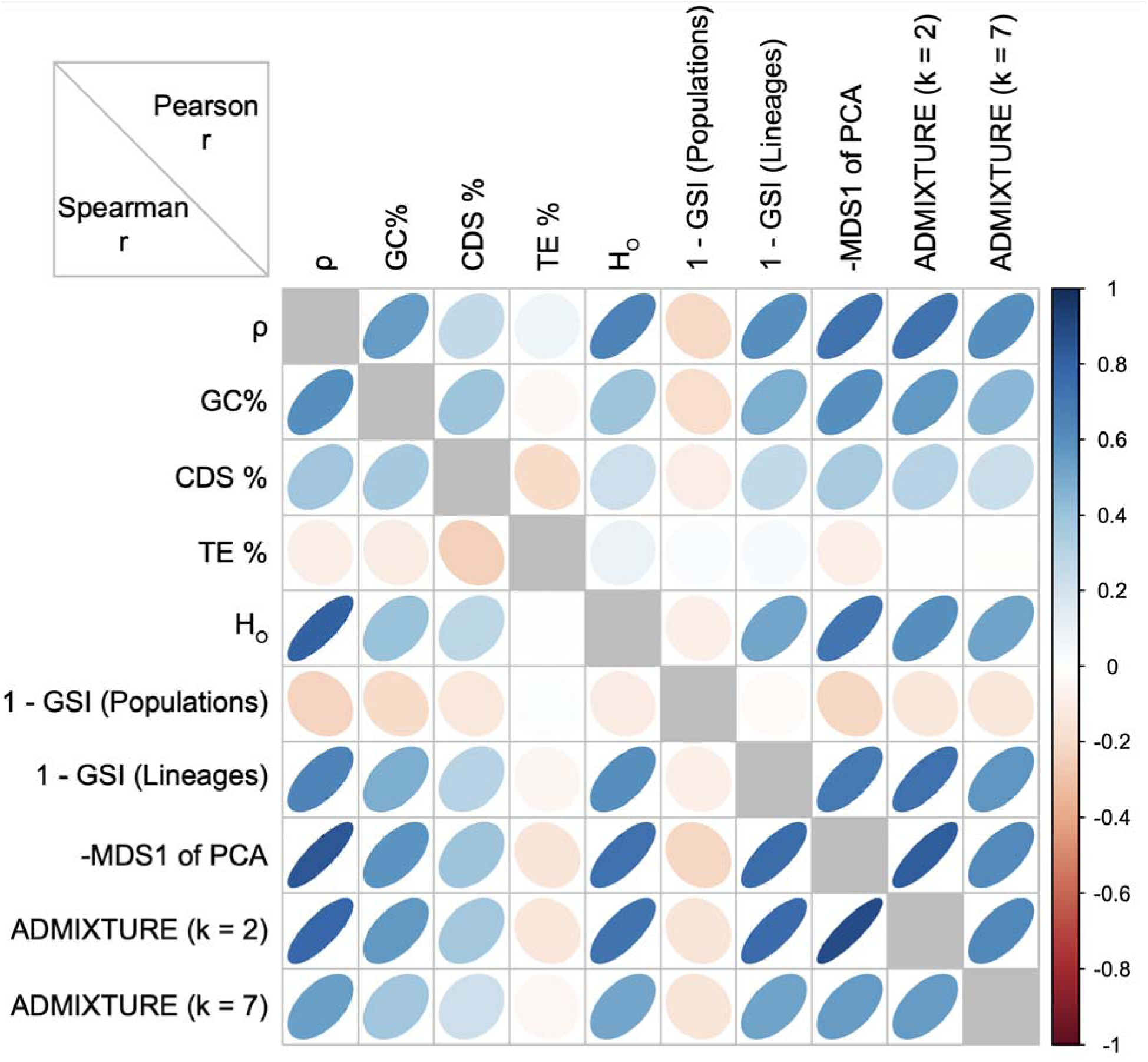
Correlation plot of genomic architecture, genomic diversity, and phylogeographic structure for 50 kbp windowed statistics. Pearson’s and Spearman’s r coefficients are plotted above and below the diagonal, respectively. Abbreviations shown in plot: mean effective recombination rate of the two main lineages (ρ); GC content (GC%); gene content (CDS%); transposable element content (TE%); mean observed heterozygosity (H_O_); genealogical sorting index at the population [1 – GSI (Pop.)] and lineage [1 – GSI (Lin.)] levels; negative multidimensional scaling axis one from principal components analyses (-MDS1); ADMIXTURE deviation relative to genome-wide analysis for two [ADMIX. (k = 2)] or seven [ADMIX. (k = 7)] assumed genetic clusters.

Phylogeographic structure patterns most different from genome-wide patterns were strongly positively correlated with recombination rate, GC content, and genetic diversity (Fig. 4; Fig. S4). In contrast, TE content had no or weak correlations with all other statistics that we calculated (Fig. 4; Fig. S4). Correlations were consistent between 50 kbp and 100 kbp estimates, with no correlations deviating more than ∼0.08 when estimated with different window sizes (Fig. S4).

Using variance partitioning, recombination rate variation most strongly explained deviations in phylogeographic structure, but interactions of recombination rate, GC content, CDS content, and TE content also explained some portions of the variance (Table 2). Most notably, genomic architecture explained approximately 60% of the variance in both (1) the MDS1 axis explaining differences in PCA-based estimates of phylogeographic structure and (2) ADMIXTURE results at the lineage level with an assumed k = 2 (Table 2).

**Table 2.**
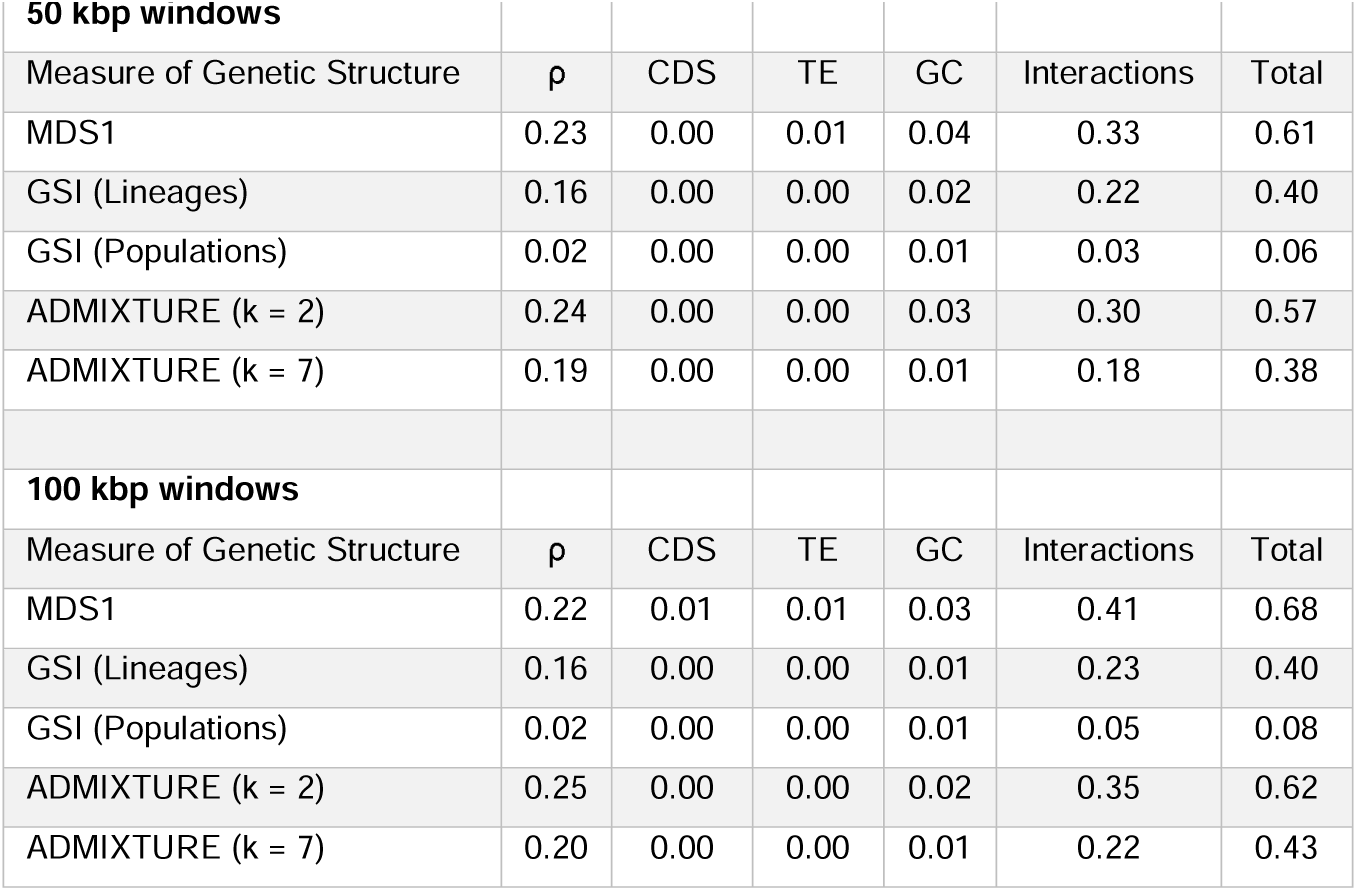
Variance partitioning where the explanatory variables are characteristics of genomic architecture and response variables are different measures of population genetic structure. Values indicative of variance proportion explained by each explanatory variable or their interactions.

The relationship between recombination rate and phylogeographic structure is exemplified when comparing the genomic windows with the highest or lowest values of recombination rate; estimates of phylogeographic structure and genetic diversity have little to no overlap in statistic distributions in high recombination regions versus low recombination regions (Fig. 5). These patterns hold regardless of whether we are looking at autosomes or the Z chromosome (Fig. 5).

**Figure 5.**
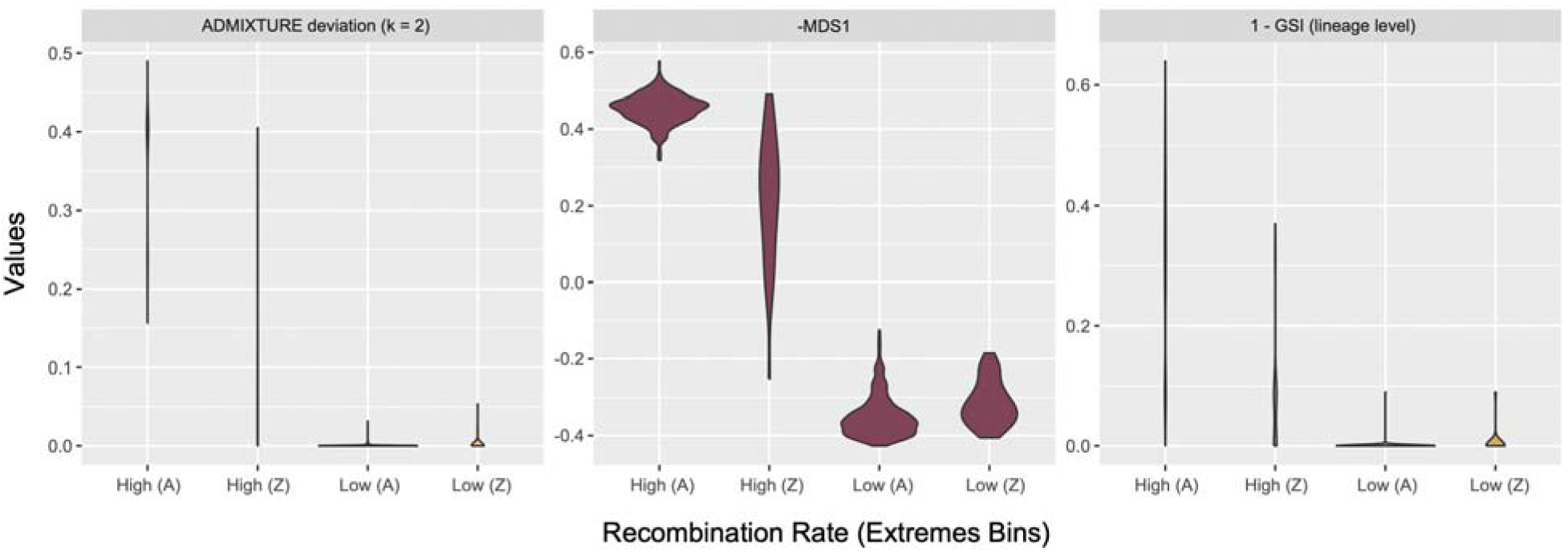
Violin plots of phylogeographic structure variation in genomic regions with very high or low recombination rates, separated for autosomal (A) and Z-linked (Z) genomic regions. 50 kbp windows with the highest or lowest 2.5% of recombination rate values are included. Higher values for ADMIXTURE deviations, 1 – GSI, and –MDS1 indicate values deviating relatively more from the genome-wide patterns of phylogeographic structure.

Because the effective population sizes for each locality varied substantially (Fig. 1; Fig. S5), we looked to see if there was a relationship between relative RTD and population size in regions with very different recombination rates. Using both the raw measurements and PICs, we found a negative association between relative rates of molecular evolution and harmonic mean effective population sizes (Fig. 6; Fig. S6). Notably, there was also a strong positive association between effective population sizes and shifts in relative rates of molecular evolution between high and low recombination regions (Fig. 6).

**Figure 6.**
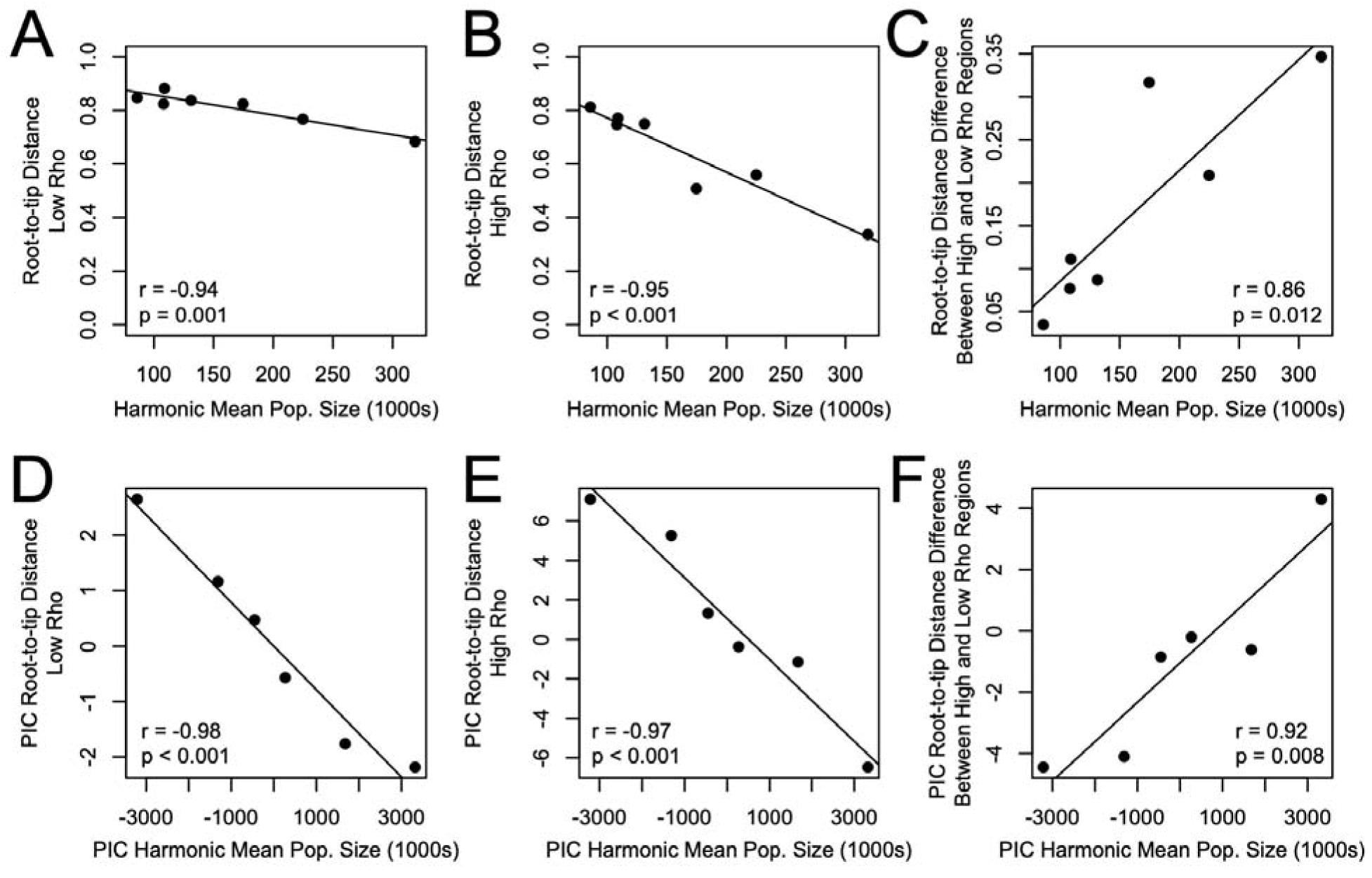
Relationship between effective population sizes and rate of evolution (measured by relative root-to-tip distance for each population) in genomic regions that vary in recombination rate. Extreme low and high recombination (rho) regions represent the quintiles of the genome with the lowest or highest recombination rates (i.e., highest 20% and lowest 20% of windows). Results for raw data are shown in (A-C) and for phylogenetic independent contrasts (PIC) in (D-F). The right panels in each row (C + F) show the shift in relative rates of evolution between high and low recombination genomic regions.

## DISCUSSION

Using whole-genome sequencing for seven populations of the Brown Creeper, we aimed to decipher how genome architecture influences phylogeographic structure variation across the genome. We found that recombination rate and other characteristics of the genome explain 6-68% of the variance in phylogeographic structure inference using different metrics and genomic sliding window sizes (Table 2).

### Phylogeographic structure shaped by genomic architecture

We used genetic clustering, ordination, and phylogenetic methods to assess phylogeographic structure in genomic sliding windows. We found that recombination rate variation was the strongest predictor of phylogeographic structure variation across the genome (Table 2). Consistent with our expectation, phylogeographic signal best representing the bifurcating evolutionary history of the Brown Creeper was identified in genomic regions with relatively low recombination rate; this was consistent across all methods used (Fig. 2). Notably, the genomic windows with the lowest recombination rates deviated little from the genome-wide patterns (Fig. 5).

Previous work in insects, birds, and mammals has shown that phylogenomic signal varies across the genome (Edelman et al., 2019; Fontaine et al., 2015; Li et al., 2019; Martin et al., 2019; Thom et al., 2024). Generally, these studies have found or suggested that the background species tree is best represented by phylogenomic patterns exhibited in genomic regions with low recombination, and that gene flow among taxa is more prevalent in genomic regions with high recombination rate (Edelman et al., 2019; Li et al., 2019; Martin et al., 2019; Thom et al., 2024). Our results in the Brown Creeper echo these findings *at a shallow evolutionary timescale*, whereby we find highest support for the genome-wide supported phylogeographic relationships in genomic regions with low recombination rate (Fig. 2; Fig. S2). We may interpret these patterns in the context of lineage sorting during the speciation process, whereby genomic regions with low introgression exhibit quicker polymorphism fixation due to relatively increased effects of linked selection and genetic drift and reduced homogenization among lineages due to gene flow.

Genomic studies investigating intraspecific population and phylogeographic structure have more commonly used PCA-based methods (e.g., LOSTRUCT) to investigate variation in genomic structure across the genome. In two species of woodpeckers, a small portion of the variance in PCA-based phylogeographic structure was found to be associated with local recombination rate (Moreira et al., 2023). We found that variation in phylogeographic structure across the genome of the Brown Creeper—inferred using both ordination and genetic clustering methods—was strongly associated with recombination rate variation (Fig. 2; Fig. S2; Table 2). We may infer that the same interactions between recombination rate variation and population genomics processes shaping variance in phylogenomic signal are also shaping variance in clustering– and ordination-based estimates of phylogeographic structure.

In contrast to generalizations about low recombination regions often reflecting a taxon’s background evolutionary history, in some scenarios we may expect regions of extremely low recombination to greatly differ from genome-wide signatures of population genetic structure. In several taxa, including sunflowers, flies, fishes, and birds, the greatest deviations from the genome-wide pattern of intraspecific population structure (inferred with PCA) have been identified in genomic regions with extremely low recombination rates that are often inferred to be polymorphic inversions within populations or species (Hale et al., 2021; Huang et al., 2020; Li and Ralph, 2019; Mérot et al., 2021; Perrier et al., 2020; Shi et al., 2021; Todesco et al., 2020; Whiting et al., 2021). In these cases, if a taxon has relatively little population structure but also exhibits polymorphic inversions, we may expect the biggest deviations in population genetic structure from genome-wide patterns to reflect these polymorphic inversions that exhibit suppressed recombination. These cases present interesting patterns to think about when interpreting genomic phylogeography data. Genomic regions with low recombination may best represent a taxon’s evolutionary history, excepting times when these regions are associated with large structural variants.

### Interaction of recombination rate, population demography, and rate of evolution

While we may expect that the interactions between recombination rate and population genomics processes will impact the rate of evolution across the genome, we did not simply observe little phylogeographic structure in genomic regions with high recombination rates and strong phylogeographic structure in genomic regions with low recombination rates. Conversely, we found strong—and differing—patterns of phylogeographic structure across the genome (Fig. 1; Fig. 3; Fig. S3). Because these differing patterns of phylogeographic structure generally separated the Central American populations and sometimes the Sierra Madre Oriental population from all others, we hypothesized that populations with relatively smaller N_E_ exhibit relatively higher evolutionary rates in genomic regions with high recombination.

Using relative RTD measures as a relative rate of molecular evolution for each population, we found that larger populations exhibited reduced rates of molecular evolution in genomic windows with high recombination rates (Fig. 6; Fig. S6) and that shifts in relative rates of molecular evolution in high versus low recombination genomic regions were positively associated with effective population sizes (Fig. 6; Fig. S6). This is consistent with simulations by Tigano and colleagues (2021) where the authors showed the interplay between recombination rate and selection causes greater genomic variance in evolutionary rates in larger populations relative to more consistent evolutionary rates across the genome in smaller populations.

We suspect that molecular evolution in genomic regions with high recombination rates is a product of complex interactions between population demography and the relative strengths of linked selection and genetic drift. In the Brown Creeper genome, recombination rate variation is positively associated with gene density (Fig. 2; Fig. 4) and we may interpret gene-dense genomic regions as targets for natural selection. Indeed, in a previous study in the Brown Creeper (Manthey et al., 2021), we showed that the two main Brown Creeper lineages exhibited relatively less neutral evolution in the gene-dense microchromosomes relative to the larger macrochromosomes. In gene-dense and high recombination regions under the effects of purifying selection, we may expect populations with relatively smaller N_E_ to accumulate substitutions faster than populations with higher N_E_ (Lanfear et al., 2014), consistent with the trends we identified here in the Brown Creeper (Fig. 6; Fig. S6).

### Implications for future phylogeographic studies

If phylogeographic structure varies widely across the genome, what are realistic and best-practice approaches for future phylogeographic work? Are studies that use few genes or reduced representation genomic datasets doomed to be wrong from the start? We expect that strong phylogeographic structure will be detected even with few genetic markers. For example, more than ten years ago we used 20 genetic markers obtained with Sanger sequencing to study the phylogeography of the Brown Creeper (Manthey et al., 2011b) and found the same general phylogeographic trends observed here with our whole-genome dataset. In contrast, any study where most of the genetic markers are taken from genomic regions where the pattern differs from a taxon’s true evolutionary history (e.g., high recombination regions in the Brown Creeper case) may run into problems inferring that true history. Additionally, in any taxa with weak population genetic structure or widespread and rampant gene flow, a whole-genome method may be the only approach to identify the few genomic regions differentiating taxa (e.g., Toews et al., 2016). Overall, we suggest using the most amount of genetic data feasible for future phylogeographic studies given sources of biological material (e.g., quality of preserved tissue) and funding available, as well as investigating the patterns and potential causes of population genetic structure variation across the genome when possible.

## CONCLUSIONS

We used whole genome sequencing to estimate how genomic architecture shapes variation in phylogeographic structure across the genome of the Brown Creeper. We found that phylogeographic structure—as measured using ordination, phylogenetic, and clustering methods—was strongly associated with regional genomic variation in recombination rates. In low recombination regions, we recovered phylogeographic structure concordant with genome-wide patterns with all three types of methods. The most divergent phylogeographic patterns were in high recombination regions; populations with small effective population sizes were distinct from all other populations due to relatively faster evolution than larger populations in these high recombination regions. Because these high recombination regions are rich in coding genes, we hypothesize that large populations have higher relative effects of purifying selection in these regions, and overall slower relative molecular evolutionary rates compared to smaller populations. Overall, our results show that phylogeographic structure may vary widely across the genome, and that effective population sizes of sampled populations and genomic architecture and their interactions will impact regional genomic variation in phylogeographic structure.

## ACKNOWLEDGEMENTS

We would like to thank the collection managers, curators, and contributors for generous tissue loans for all individuals used in this study from the following museums: Burke Museum of Natural History and Culture, and University of Kansas Biodiversity Institute. Sequencing was supported by Texas Tech University start-up funding to JDM and a generous donation from the Ferguson family to GMS and the Denver Museum of Nature and Science. Support was also provided by NSF awards #1953688 to JDM and #0814841 to GMS. The High-Performance Computing Center at TTU supported computational analyses.

## DATA ACCESSIBILITY

All raw resequencing data uploaded to NCBI’s SRA database under BioProject PRJNA1068991. All code used for analysis on this project is available on GitHub: (<u>github.com/jdmanthey/certhia_phylogeography</u>).

## AUTHOR CONTRIBUTIONS

JDM and GMS designed the study. JDM performed laboratory and bioinformatic work and wrote the first draft of the manuscript. JDM and GMS contributed to revising and improving the final draft of the manuscript.

## DECLARATION OF INTEREST

The authors declare no conflicts of interest.

## SUPPLEMENTAL TABLES AND FIGURES

**Table S1.** Sampling metadata and characteristics of sequencing amount and coverage when aligned to the reference genome. Samples from Utah, Morelos, Jalisco, and the outgroup were retrieved from the NCBI SRA and originally sequenced for a previous study by Manthey and colleagues (2021).

| Museum # | Lineage | Locality | Lat. | Long. | Raw bp | Filtered bp | Coverage |
| --- | --- | --- | --- | --- | --- | --- | --- |
| UWBM103417 | South | Copan, HN | 14.866 | -89.050 | 40,501,485,458 | 34,714,897,849 | 22.79 |
| UWBM105618 | South | Copan, HN | 14.866 | -89.050 | 30,149,924,172 | 26,420,616,801 | 18.55 |
| UWBM93696 | South | Copan, HN | 14.866 | -89.050 | 40,591,948,048 | 36,089,400,965 | 26.06 |
| UWBM113447 | South | Chiapas, MX | 16.691 | -92.606 | 37,741,499,908 | 33,063,229,986 | 23.32 |
| UWBM113461 | South | Chiapas, MX | 16.691 | -92.606 | 32,696,721,580 | 27,488,157,228 | 18.51 |
| UWBM113462 | South | Chiapas, MX | 16.691 | -92.606 | 48,519,014,074 | 42,841,474,607 | 27.09 |
| UWBM106936 | South | Morelos, MX | 19.0873 | -99.194 | 40,136,994,108 | 35,563,542,819 | 26.35 |
| UWBM107890 | South | Jalisco, MX | 21.882 | -103.865 | 43,601,103,228 | 38,563,138,625 | 26.14 |
| UWBM107897 | South | Jalisco, MX | 21.882 | -103.865 | 30,536,607,690 | 27,038,872,945 | 18.55 |
| UWBM106768 | South | Nuevo Leon, MX | 24.872 | -100.224 | 42,330,853,397 | 37,109,282,807 | 27.62 |
| UWBM111691 | South | Nuevo Leon, MX | 24.872 | -100.224 | 41,523,724,956 | 36,804,470,356 | 26.21 |
| UWBM111708 | South | Nuevo Leon, MX | 24.872 | -100.224 | 43,392,483,440 | 36,820,385,840 | 27.04 |
| UWBM113198 | North | California, US | 36.245 | -121.700 | 40,117,812,880 | 35,379,027,102 | 26.26 |
| UWBM112794 | North | California, US | 36.245 | -121.553 | 37,692,762,846 | 33,128,089,974 | 23.33 |
| UWBM112795 | North | California, US | 36.245 | -121.553 | 44,376,620,806 | 38,809,177,747 | 28.71 |
| UWBM113162 | North | Utah, US | 37.317 | -113.455 | 40,934,172,636 | 36,217,919,293 | 26.25 |
| UWBM113168 | North | Utah, US | 37.317 | -113.455 | 39,295,062,200 | 34,925,676,137 | 25.72 |
| UWBM113167 | North | Utah, US | 37.377 | -113.467 | 42,535,860,138 | 37,425,027,332 | 27.55 |
| UWBM107061 | North | West Virginia, US | 38.289 | -79.937 | 43,886,484,470 | 38,699,509,454 | 28.65 |
| UWBM112055 | North | West Virginia, US | 38.289 | -79.937 | 47,972,794,224 | 40,458,960,409 | 29.36 |
| UWBM112054 | North | West Virginia, US | 38.641 | -79.838 | 39,431,412,482 | 34,774,499,050 | 25.96 |
| KU92846 | Outgroup | England |  |  | 21,417,629,506 | 19,195,682,239 | 13.58 |

**Figure S1.**
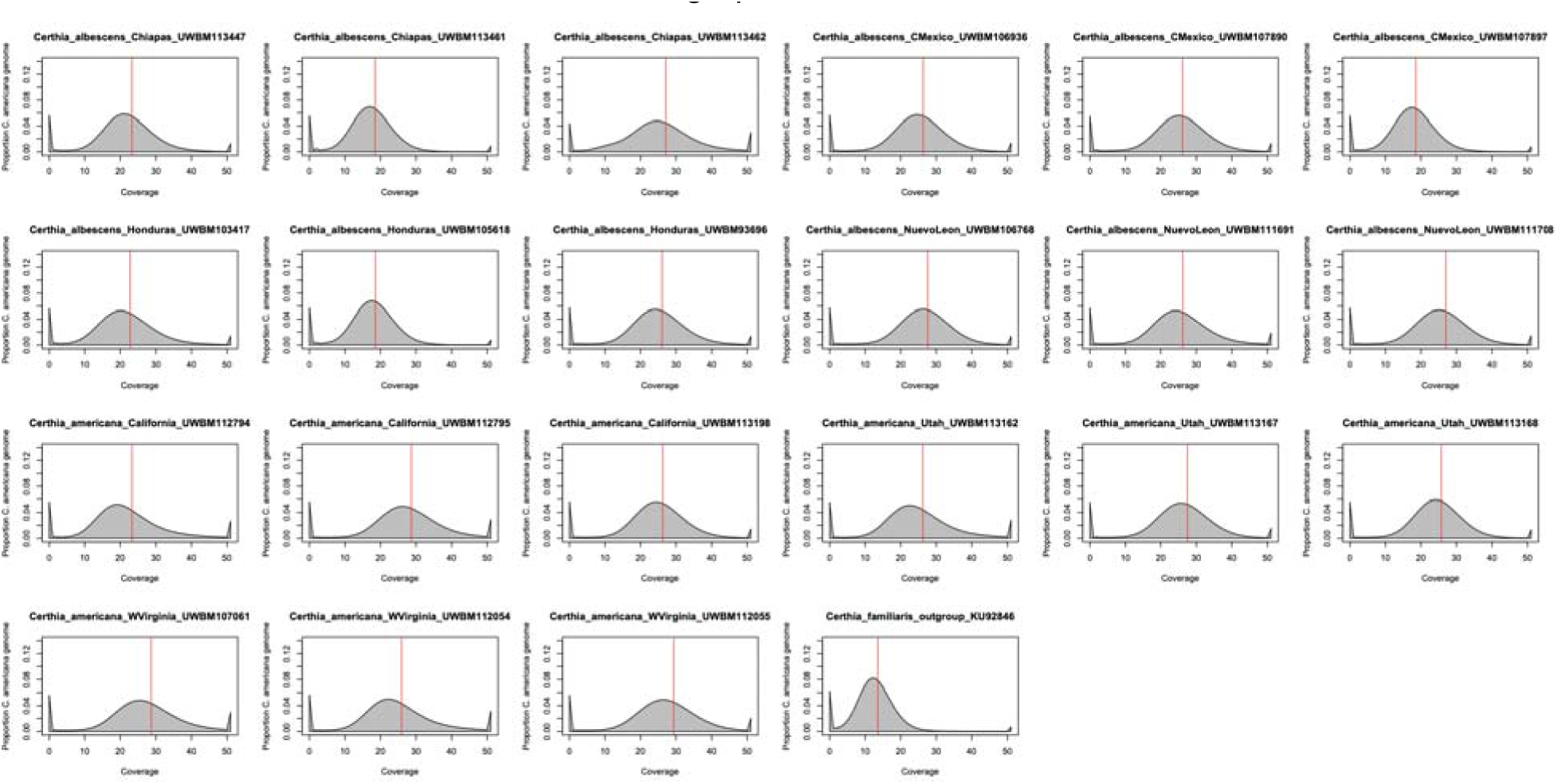
Sequencing coverage aligned to the reference genome for each individual. The vertical red lines indicate the mean coverage per individual.

**Figure S2.**
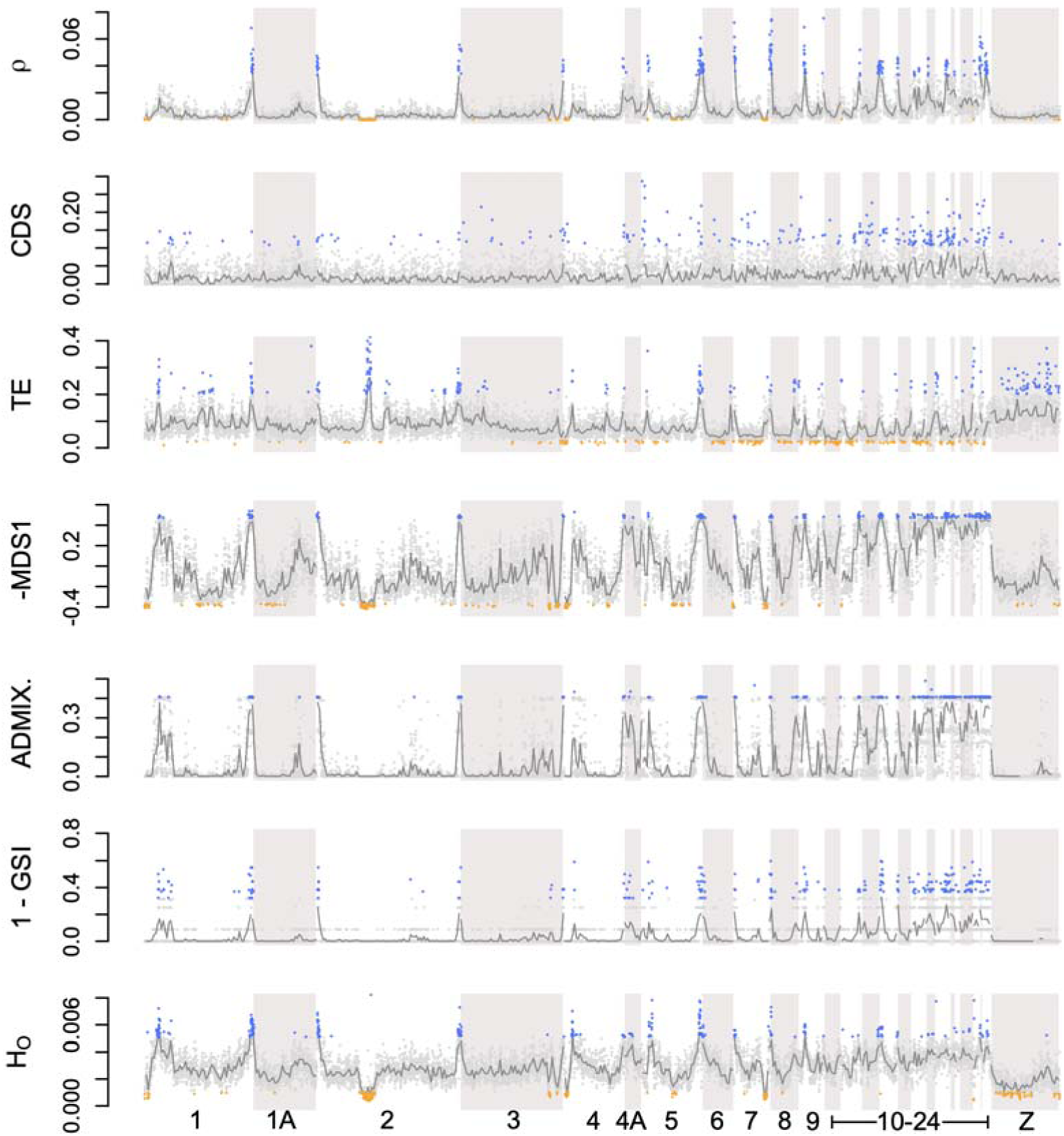
Genomic architecture and phylogeographic structure variation across the genome in 100kbp sliding windows. Gray points indicate window values and dark gray lines indicate means across ten windows (i.e., 1 Mbp windows). Blue indicates top 2.5% of outliers and orange indicates bottom 2.5% of outliers for each statistic. Bottom outliers not shown for CDS, ADMIX., and GSI as these each have large proportions of values equal to zero. Abbreviations: mean effective recombination rate of the two main lineages (ρ), gene content (CDS), transposable element and repetitive DNA content (TE), multidimensional scaling axis one from principal components analyses (-MDS1), mean genealogical sorting index for the two main *Certhia* lineages (1 – GSI), ADMIXTURE deviation relative to genome-wide analysis (ADMIX.), and mean genetic diversity across individuals as measured using observed heterozygosity (H_O_). MDS1 is plotted as negative and GSI is plotted as (1 – GSI) so that deviations from expected phylogeographic structure for MDS1, GSI, and ADMIXTURE are all represented as higher values.

**Figure S3.**
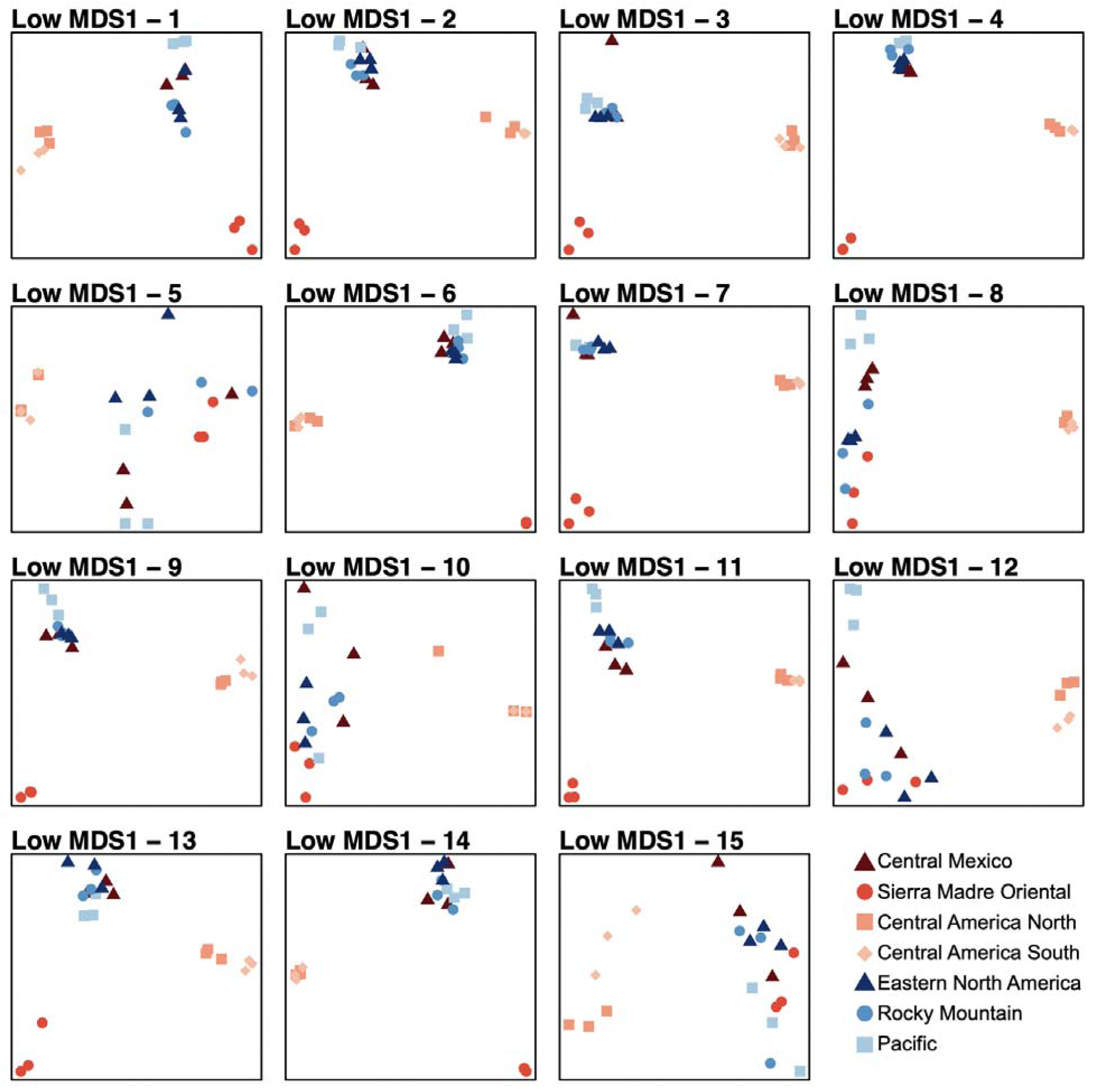
The 15 most extreme outlier principal components analysis (PCA) windows identified from those windows with low MDS dimension one values.

**Figure S4.**
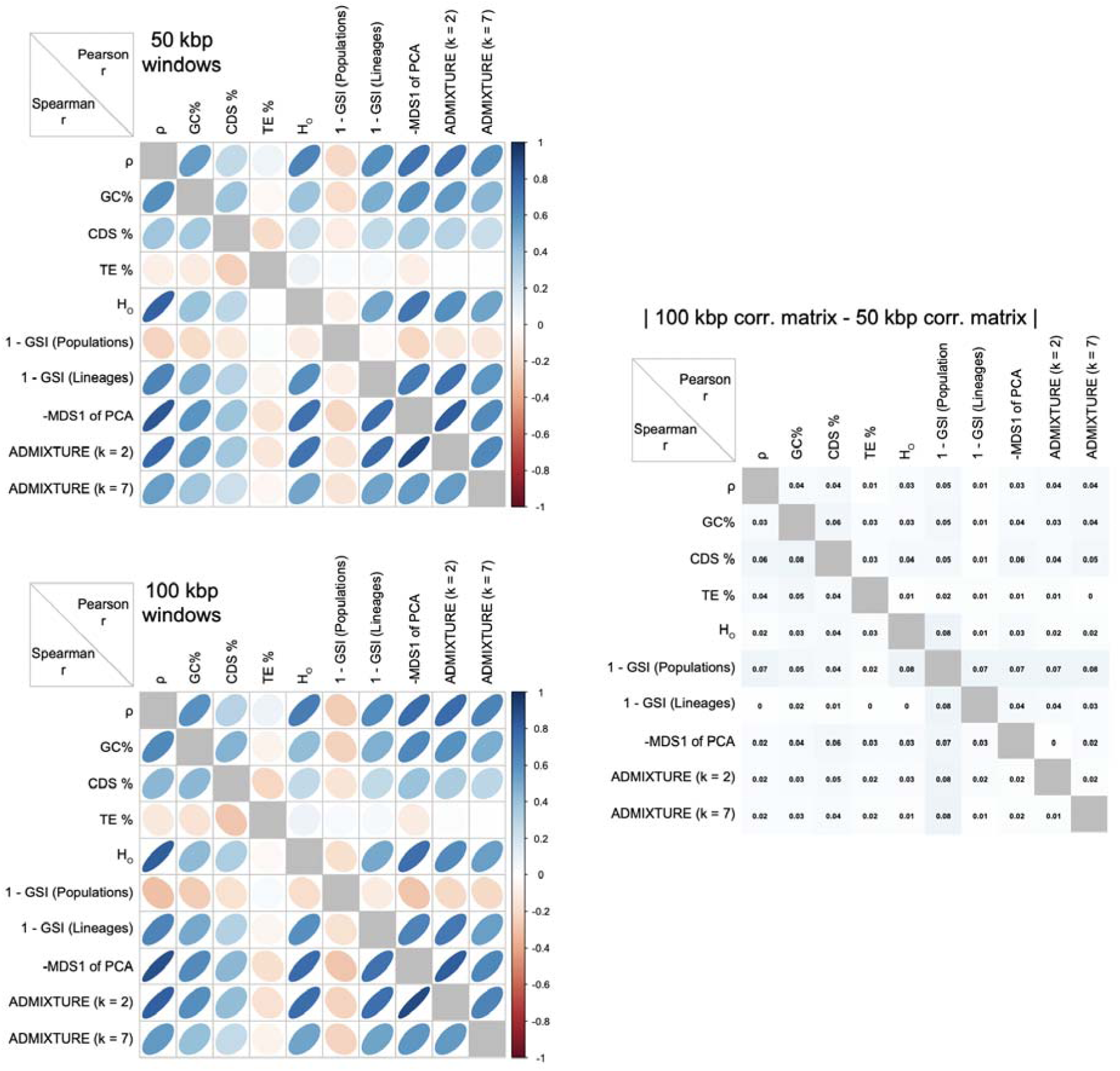
Correlation plot of genomic architecture, genomic diversity, and phylogeographic structure for (A) 50 kbp and (B) 100 kbp windowed statistics. Pearson’s and Spearman’s r coefficients are plotted above and below the diagonal, respectively. In (C), the absolute value difference between correlations in 50 kbp and 100 kbp windows. Abbreviations shown in plot: mean effective recombination rate of the two main lineages (ρ); GC content (GC%); gene content (CDS%); transposable element content (TE%); mean observed heterozygosity (H_O_); genealogical sorting index at the population [1 – GSI (Pop.)] and lineage [1 – GSI (Lin.)] levels; negative multidimensional scaling axis one from principal components analyses (-MDS1); ADMIXTURE deviation relative to genome-wide analysis for two [ADMIX. (k = 2)] or seven [ADMIX. (k = 7)] assumed genetic clusters.

**Figure S5.**
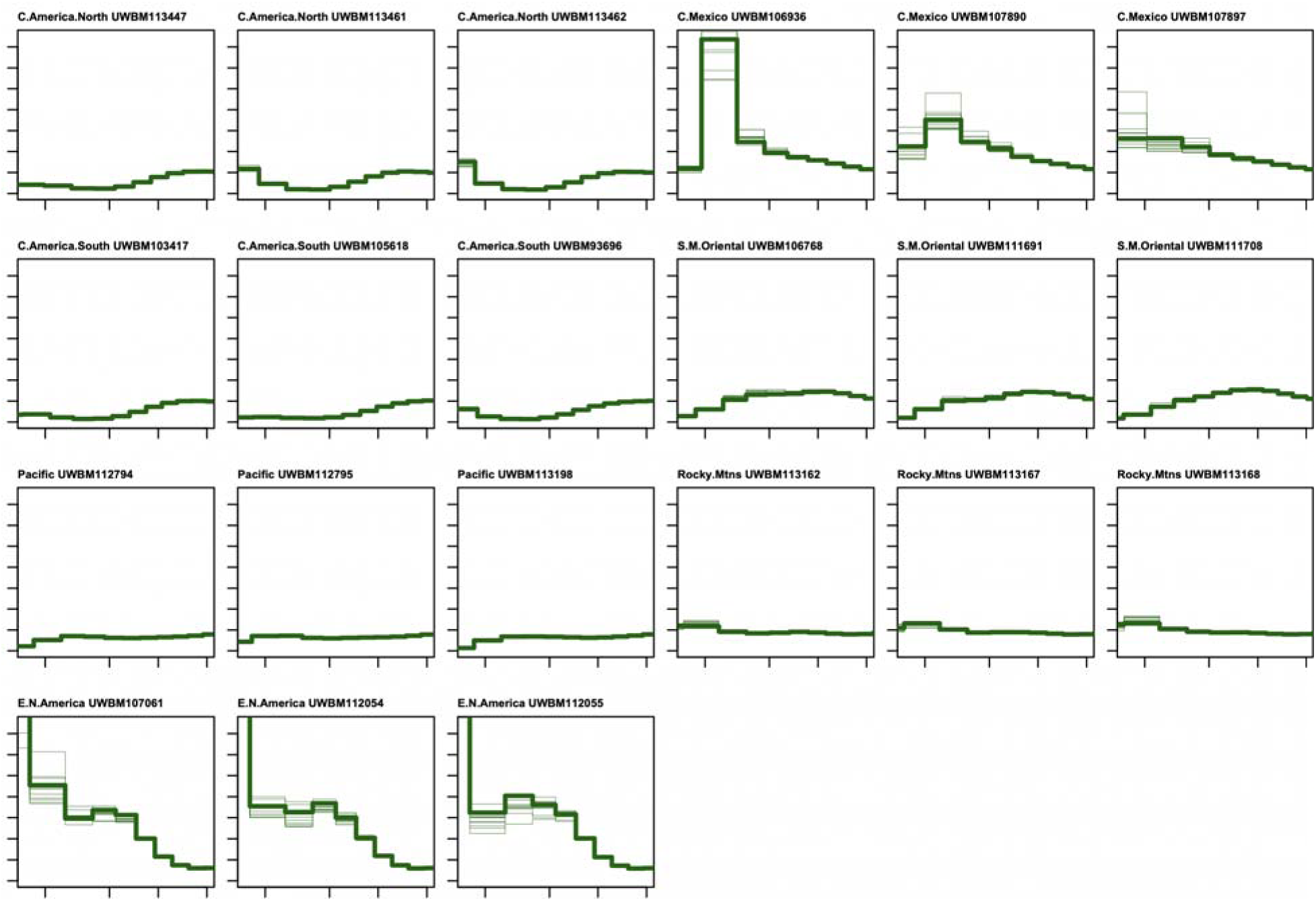
Demographic history estimated with MSMC2 for each individual. Thick lines indicate the whole genome estimate and thin lines indicate results from ten bootstrap replicates. Sample numbers correspond with those reported in Table S1.

**Figure S6.**
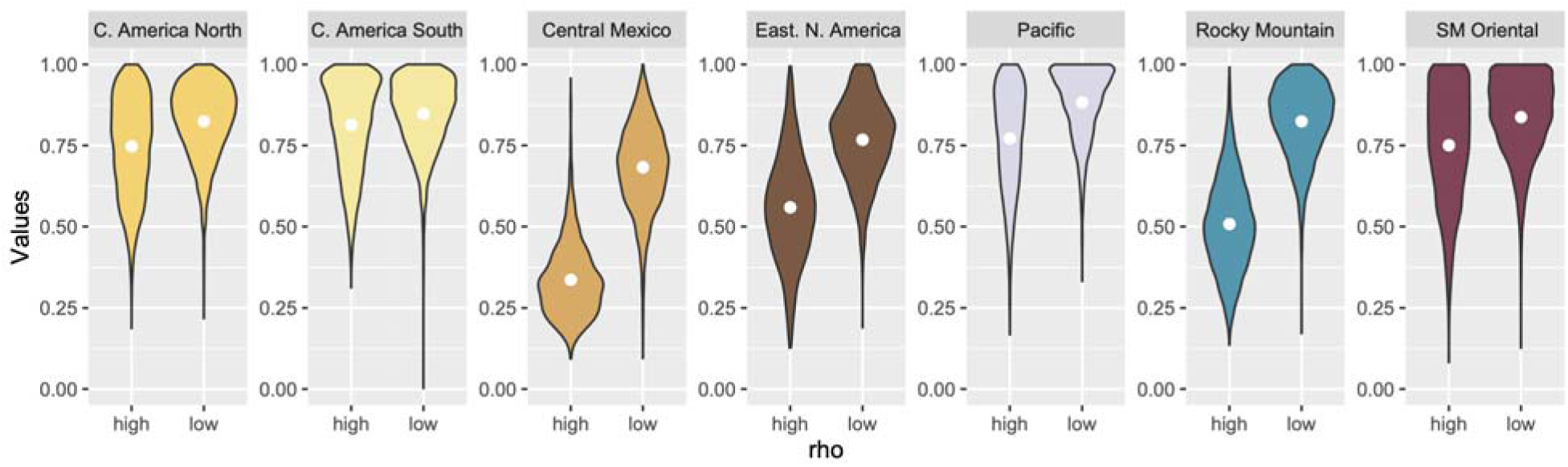
Violin plots of evolutionary rates (as measured by relative root-to-tip distances) for each population in the upper and lower genomic quintiles of high or low recombination (rho) values.

## Notes

### Competing Interest Statement

The authors have declared no competing interest.

### Summary of Updates

Aspects of the intro, methods, results, and discussion have been reworked. Some of the figures have been modified to reflect some changes to methods/results.

